# The gut parasite *Nosema ceranae* impairs olfactory learning in bumblebees

**DOI:** 10.1101/2021.05.04.442599

**Authors:** Tamara Gómez-Moracho, Tristan Durand, Mathieu Lihoreau

**Affiliations:** Research Center on Animal Cognition (CRCA), Center for Integrative Biology (CBI); CNRS, University Paul Sabatier, Toulouse, France

**Keywords:** Bumble bees, *Bombus terrestris*, learning and memory, PER

## Abstract

Pollinators are exposed to numerous parasites and pathogens when foraging on flowers. These biological stressors may affect critical cognitive abilities required for foraging. Here, we tested whether exposure to *Nosema ceranae*, one of the most widespread parasite of honey bees also found in wild pollinators, impacts cognition in bumblebees. We investigated different forms of olfactory learning and memory using conditioning of the proboscis extension reflex. Seven days after feeding parasite spores, bumblebees showed lower performance in absolute and differential learning, and reversal learning than controls. Long-term memory was also slightly reduced. The consistent effect of *N. ceranae* exposure across different types of olfactory learning indicates that its action was not specific to particular brain areas or neural processes. We discuss the potential mechanisms by which *N. ceranae* impairs bumblebee cognition and the broader consequences for populations of pollinators.

## 1. Introduction

Pollinators, such as bees, rely on a rich cognitive repertoire to collect pollen and nectar on flowers. These include associative learning and memories of floral traits like odours, shapes, colours, and textures, to identify profitable resources (Giurfa, 2015; Menzel, 2012), and spatial cues to navigate (Collett et al., 2013). Any disruption of these cognitive abilities by environmental stressors can considerably reduce the foraging performances of bees, ultimately compromising brood development and survival (Klein et al., 2017).

In particular, foraging bees are exposed to a number of parasites that can affect their physiology and behaviour (Gómez-Moracho et al., 2017). The microsporidia *Nosema ceranae* is one of the most prevalent parasites of bees worldwide with a large range of hosts including honey bees (Higes et al., 2006), bumblebees (Plischuk et al., 2009), solitary bees (Ravoet et al., 2014), but also other flower visitors like wasps (Porrini et al., 2017). Insects get infected by ingesting parasite spores from contaminated water or pollen (Higes et al., 2008), or during physical contacts with contaminated individuals (Smith, 2012). The spores invade the gut epithelial cells of the bee where they develop (Holt et al., 2013). *Nosema* degenerates the gut epithelium of the host (Higes et al., 2007), alters its metabolism (Mayack and Naug, 2009) and disrupts the immune response (Antúnez et al., 2009). In honey bees, this causes a disease (nosemosis) believed to contribute to colony collapse disorder (Cox-Foster et al., 2007).

Infected honey bees also show impaired navigation (Wolf et al., 2014) and increased flight activity (Dussaubat et al., 2013) suggesting their cognitive abilities are affected. Few studies have explored this possibility and their results so far are mixed (Bell et al., 2020; Charbonneau et al., 2016; Gage et al., 2018; Piiroinen et al., 2016; Piiroinen and Goulson, 2016). Most of them were conducted on honey bees using Pavlovian olfactory conditionings of the proboscis extension reflex (PER) in which harnessed bees are trained to associate an odour, or a combination of odours, with a sucrose reward (Takeda, 1961). Honey bees showed reduced absolute learning (when an odour is paired to a reward) between days 7 and 23 after parasite exposure ((Bell et al., 2020; Gage et al., 2018; Piiroinen and Goulson, 2016); but see (Charbonneau et al., 2016)). Short-term memory was impaired after 7 and 15 days (Gage et al., 2018), but not after 9 (Piiroinen and Goulson, 2016) or 23 days (Bell *et al*., 2020). Long-term memory was affected after 7 days but not after 15 days (Gage et al., 2018). Only two studies explored these effects on bumblebees. One suggests a slight impairment of absolute learning (Piiroinen and Goulson, 2016), and both report an absence of effect on memory (Piiroinen et al., 2016, Piiroinen and Goulson 2016). Note however that in these two studies less than 3% of the bumblebees exposed to the parasite showed signs of infection after the behavioural tests (i.e. PCR positive).

Given the expanding geographical distribution of *N. ceranae* worldwide (Klee et al., 2007) and its prevalence in domesticated and wild bees (Plischuk et al., 2009; Porrini et al., 2017; Ravoet et al., 2014), clarifying its influence on host cognition has become an urgent matter. In particular, other critical forms of learning, such as the ability to associate one of two odours with a reward (differential learning) and reverse this association (reversal learning) have so far been unexplored. These types of learning are essential to discriminate flowers. They involve different brain centers and may thus be more or less impacted by the parasite if its effect is specific (Boitard et al., 2015; Giurfa and Sandoz, 2012).

Here we build on a recently established method yielding high the rates of experimental infection by *N. ceranae* (Gómez-Moracho et al., 2021) to study the impact of the parasite on different cognitive tasks in bumblebees. We used PER conditioning to compare the olfactory learning and memory performances of control bumblebees, bumblebees exposed to the parasite, and bumblebees contaminated by the parasites (PCR positive) at seven days post exposure.

## 2. Material and methods

### 2.1. Bumblebees

We used bumblebees (*B. terrestris*) from 14 commercial colonies acquired from Biobest (Westerlo, Belgium). Before the experiments, we verified the absence of *N. ceranae* (Martín-Hernández *et al*., 2007), and other common parasites (*N. bombi* (Klee *et al*., 2006); *Crithidia bombi* (Schmid-Hempel and Tognazzo, 2010)) in a PCR using 15 bumblebees from each colony. We maintained bumblebees in their original colonies with *ad libitum* access to the syrup provided by the manufacturer and germ-free pollen (honey bee collected pollen exposed to UV light for 12 hours), in a room at 25±1°C under a 12 h light: 12 h dark photocycle.

### 2.2. *N. ceranae* spores

We obtained fresh spores from naturally infected honey bee colonies (*Apis mellifera*) maintained at our experimental apiary (University Toulouse III, France). To prepare spore solutions, we dissected the gut of 15 honey bees and crushed them in 15 mL of distilled H_2_O. We confirmed by PCR the presence of *N. ceranae* and the absence of *N. apis* (another common parasite of honey bees) in each homogenate (Martín-Hernández et al., 2007), and purified them following standard protocols (Fries et al., 2013). We centrifuged homogenates in aliquots of 1 mL at 5,000 rpm for 5 minutes and re-suspended the pellet in 500 μL of dH_2_O by vortexing. This was repeated three times to obtain a spore solution of 85% purity (Fries et al., 2013). We counted *N. ceranae* spores in an improved Neubauer haemocytometer (Cantwell, 1970) in a light microscope (x400) and adjusted the spore inoculum to 15,000 spores/μL in 20% (w/w) of sucrose solution. Spore solutions were used within the same week they were purified.

### 2.3. Parasite exposure and experimental conditions

We exposed bumblebees to *N. ceranae* as described in Gómez-Moracho *et al*., (2021). Briefly, we confined individual bumblebees in a Petri dish during 5 h without food. We then exposed some bumblebees to a 20 μL drop of inoculum containing 300,000 *N. ceranae* spores. Control bumblebees received 20 μL of sucrose solution (20% w/w). We only used bumblebees that consumed the entire drop of sucrose within the next 2 h. We then allocated bumblebees into microcolonies of 20-25 individuals, containing a gravity feeder with *ad libitum* access to food (Kraus et al., 2019). Since diet can affect host-parasites relationships (Frost et al., 2008) we provided bumblebees an artificial diet with a protein to carbohydrates ratio of 1:207 previously shown to elicit highest *N. ceranae* prevalence in bumblebees (Gómez-Moracho et al., 2021). The diet was made with a fixed total amount of nutrients of 170 g/L (protein + carbohydrates) and 0.5% of vitamin mixture for insects (Sigma, Germany). Carbohydrates were supplied as sucrose (Euromedex, France). Proteins consisted in a mixture of casein and whey (4:1) (Nutrimuscle, Belgium) (Gómez-Moracho et al., 2021). We kept bumblebee microcolonies in a room at 25±1 °C with a 12 h light: 12 h dark photoperiod until the behavioural tests. Every day, we renewed the diet and removed dead bumblebees.

### 2.4. Behavioural experiments

We tested the cognitive performances of bumblebees using PER at day 7 after parasite exposure. The day before the behavioural tests we kept diet to low levels (~200 μL/bumblebee) to keep bumblebees motivated for PER experiments. Three hours before the behavioural tests we collected bumblebees, chilled them in ice for 5 min and restrained them in a modified 2 mL Eppendorf tube (hereafter, capsule) that we cut in length to fit each bumblebee (adapted from Toda *et al*., (2009); Figure 1A). Bumblebees were tested in the horizontal position and could slightly move inside the tube. We found these conditions better suited to perform PER experiments with bumblebees than the classical vertical harnessing used for honey bees (Giurfa and Sandoz, 2012), in which bumblebees appeared paralysed (unpublished data). Once in the capsule, we kept the bumblebees in the dark, in an incubator at 28°C, with no access to food. All bumblebees that finished the conditioning protocols were kept at −20°C for later analyses of their infection status through PCR.

**Figure 1.**
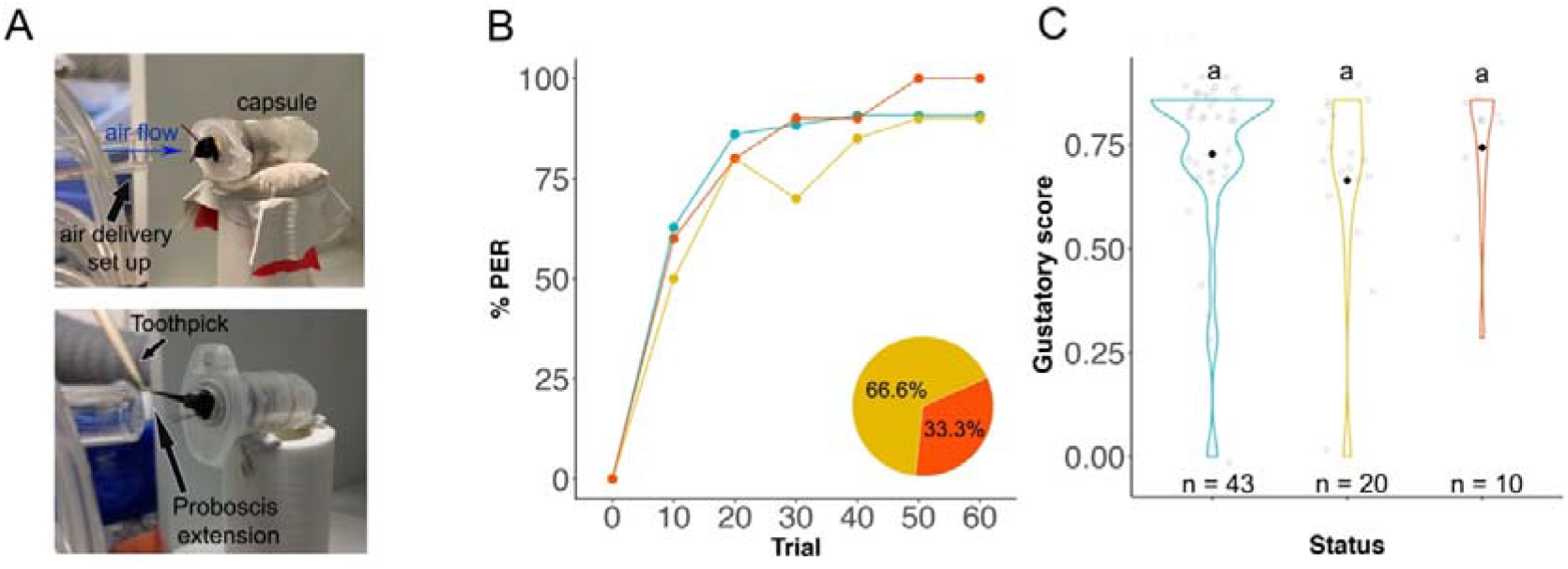
Odour delivery set up and sucrose responsiveness. **A)** Photo of a bumblebee during odour conditioning. The bumblebee is placed inside the capsule in front of the air delivery set up. After the odour is delivered, sucrose is presented with a toothpick to the bumblebees which extends its proboscis to drink the reward. **B)** Proportion of control (blue), exposed negative (yellow) and exposed positive (red) bumblebees responding to an increase gradient of sucrose concentrations. Pie chart represents the percentage of exposed negative and exposed positive bees (n total=30). **C)** Violin plots show gustatory score of bumblebees as the sum of all responses for each bumblebee. Black diamonds represent the mean score for each infection status. White dots represent the score of each individual. n is the sample size. Letters above violin plots represent significant differences between status (GLMM; p < 0.05). n is the sample size.

### 2.5. Sucrose responsiveness

We tested the sucrose responsiveness of bumblebees to control for potential influences of *N. ceranae* on their reward perception or feeding motivation. We presented seven sucrose solutions to each bumblebee, from concentrations of 0% (pure water) to 60% (w/w), with increments of 10% (Graystock et al., 2013). For each concentration, we touched the antennae of the bumblebee with a toothpick soaked in the corresponding sucrose solution to elicit PER. We presented solutions in an increasing concentration gradient with an inter-trial interval of 5 minutes between concentrations. We scored a positive response if bumblebees extended their proboscis after a solution presentation. We discarded bumblebees responding to water (i.e. 0% sucrose solution) to avoid the effect of thirst on sucrose responsiveness (Baracchi et al., 2018). The sum of the number of PER responses divided by the number of concentration tested formed the individual gustatory score (Scheiner et al., 2013).

### 2.6. Conditioning experiments

All experiments shared the same general protocol (Figure S1). An encapsulated bumblebee (Figure 1A) was placed 1.5 cm ahead of an automated conditioning setup (described in (Aguiar et al., 2018)) delivering a continuous stream of odourless air at 1.2 ml/s to which specific odours were selectively added (Raiser et al., 2017). We used two odorants as conditioned stimulus (CS): nonanal and phenylacetaldehyde (Palottini et al., 2018), in a 1:100 dilution in mineral oil. Before conditioning, we tested the responsiveness of bees to sucrose by touching both antennae with a toothpick soaked in 50% (w/w) sucrose solution without allowing them to lick. Bumblebees extending their proboscis were considered motivated and kept for the experiments. Conditioning trials (Figure S1c) consisted in 15 seconds of odourless airflow, followed by 6 seconds of CS, and 3 seconds of unconditioned stimulus (US) (i.e. 50% sucrose solution applied with a toothpick on the bumblebee’s antennae), with 2 seconds of overlap between CS+US, and 20 seconds of odourless airflow (Aguiar et al., 2018). The inter-trial interval was 10 minutes. An air extractor was placed behind the bumblebee to prevent odorant accumulation during CS delivery. Bumblebees extending their proboscis within 3 seconds of US presentation (i.e. 2s CS+US and 1s US) were allowed to lick the toothpick soaked in sucrose (50%, w/w). In unrewarded trials (see reversal learning and memory tests) no US was applied. We scored a conditioned response if the bumblebee extended its proboscis to the odour delivery before sucrose presentation (Figure S1c). Bumblebees that responded to the odour in the first conditioning trial were discarded from the analyses. We used conditioned responses to calculate three individual scores for each bumblebee, describing its performance during conditioning (i.e. acquisition score), at the end of conditioning (i.e. learning score) or during memory retrieval (i.e. memory score) (Monchanin et al., 2020; see details below). Exposed and control bumblebees were always conditioned in parallel.

#### 2.6.1. Absolute learning, short-term memory and long-term memory

We tested the effects of *N. ceranae* on the ability of bumblebees to associate an odour with a reward. This form of learning only requires peripheral brain centers, i.e. the antennal lobes (Giurfa and Sandoz, 2012). We trained bumblebees in a spaced 3-trial absolute conditioning learning (Figure S1a) that was shown to generate robust long-term memory in bees (Menzel et al., 2001). We used the same rewarded odour (CS+) during training of a given bumblebee, but both nonanal and phenylacetaldehyde were used as a CS+ for different bumblebees. Only bumblebees that responded to the US in the three trials (i.e. efficient bumblebees) were kept for the analyses (Table S2). For each of these bees we calculated an acquisition score (sum of PER responses (i.e. 0-2) divided by the number of trials (i.e. 3)), and a learning score (1 if they responded to CS+ in the last trial or 0).

We tested memory retention in efficient bumblebees that made at least one conditioned response in either one of the last two trials. We performed tests either 1 h (i.e. short-term memory; STM) or 24 h (i.e. long-term memory; LTM) after the last acquisition trial. For tests performed after 24 h, bumblebees were fed until satiation with 50% (w/w) sucrose solution right after conditioning. In bees, LTM is dependent on protein synthesis whereas STM is not (Menzel, 2001). Studying these two types of memories was thus a mean to explore whether exposure to *N. ceranae* interfered with protein synthesis. We presented bumblebees the two odorants without any reward: the odour used as a CS+ to test for memory formation, and the second odour as a novel odorant (NOd), to control for potential generalization (Villar et al., 2020). For example, when nonanal was used as CS+, phenylacetaldehyde was used as a NOd, and *vice versa*. Memory formation was considered when bumblebees extended their proboscis in response to the CS+ but not to the NOd, and thus had a memory score of 1 (0 otherwise). Just after the memory test, we tested the motivation of bumblebees by touching their antennae with a toothpick soaked with 50% (w/w) sucrose solution. Bumblebees that did not respond to the US were discarded for the analyses (i.e. 14.06% in LTM; Table S2).

#### 2.6.2. Reversal learning

We tested the effects of *N. ceranae* on the ability of bumblebees to learn to discriminate two odours and reverse the task. This form of learning involves a first phase of differential conditioning that requires the antennal lobes but is not dependant on central brain centers, i.e. mushroom bodies, and a second phase of reversal learning that requires both the antennal lobes and the mushroom bodies (Boitard et al., 2015). In the differential learning phase (phase 1), we trained bumblebees to discriminate between two odours. This phase consisted in 10-trial blocks (Figure S1b), five with each odour that was either paired with the US (A+) or unpaired (B-), presented in a pseudo-random order. The rewarded and unrewarded odours were randomized on different training days. Bumblebees that did not respond to US in two or more trials were discarded for the analyses and for reversal phase. In the reversal learning phase (phase 2) we trained bumblebees to invert the first learnt contingency. The reversal phase started 1h after the end of the differential phase. Here we trained bumblebees in 12 trial-blocks, six with each odour presented in a pseudo-random order. The previously rewarded odour was not associated with a reward anymore (A-) while the previously unrewarded odour became rewarded (B+). To start with the same level of learning, we analysed reversal learning in bumblebees that responded to A-either in the first and/or second presentation of this odour during the reversal phase (Table S3). We analysed the performance of bumblebees in each phase separately by attributing them an acquisition score (sum of all trials where the bee responded to CS+ but not to CS-, divided by the number of trials) and a learning score (1 if the bumblebee responded to CS+ but not to CS-in the last trial) for each phase.

### 2.7. Infection status

We assessed the infection status of bumblebees that finished the tests in a PCR using the primers 218MITOC (Martín-Hernández et al., 2007). For each bumblebee, the entire gut was extracted, homogenised in sterile dH2O and vortexed with 2 mm glass beads (Labbox Labware, Spain). Genomic DNA was extracted using Proteinase K (20 mg/mL; Euromedex, France) and 1 mM of Tris-EDTA Buffer (pH = 8). A sample with *N. ceranae* spores was included in each round of extraction as positive control. PCRs were performed with the Taq Polymerase Direct Loading Buffer (5 U/μL; MP Biomedicals, CA) following manufacturer’s instructions. We used a final volume of 25 μL with 0.4 μM of each pair of primers (Martín-Hernández et al., 2007), 200 μM of dNTPs (Jena Biosciences, Germany), 0.48 μg/μL of BSA (Sigma, Germany) and 2.5 μL of DNA sample. PCR reactions were carried out in a S1000™ Thermal Cycler (Biorad, CA). Thermal conditions were 94 °C for 2 min, 35 cycles of 94 °C for 30 s, 61.8°C for 45 s and 72 °C for 2 min, and a final step of 72 °C for 7 min. The length of PCR products (i.e. 218 pb) was checked in a 1.2% agarose gel electrophoresis stained with SYBR Safe DNA Stain (Edvotek, Washington DC). Positive and negative controls of PCR were run in parallel. Based on the PCR results we classified bumblebees in three different infection statuses, i.e. control, exposed negative or exposed positive. Bumblebees that were not exposed to the parasite but nevertheless showed a positive result in PCR were excluded from the analyses (i.e. 6.26%; 23 out of 367 control bumblebees; Tables S1, S2 and S3).

### 2.8. Statistical analyses

All analyses were conducted in RStudio (version 1.4.1106). We evaluated the effect of parasite exposure and infection on gustatory, acquisition, learning and memory scores. The gustatory scores and the acquisition scores (i.e. absolute and reversal conditioning) were standardized and analysed in a Generalized Linear Mixed Model (GLMM) (package [glmmTMB]; (Brooks et al., 2017)). The response to the first and last sucrose concentrations, as well as the learning and memory scores were analysed in a binomial GLMM (package [lme4]; (Bates et al., 2015)). In all models, bee infection status was used as a fixed factor, while bee identity and odorant were used as random factors. We performed Tukey post-hoc pairwise comparisons (package [multcomp]; (Hothorn et al., 2008)) to assess the relationship between the three bee statuses. Raw data are available online (doi:10.5281/zenodo.4376362).

## 3. Results

### 3.1. Parasite exposure did not influence sucrose responsiveness

We tested responsiveness to different sucrose concentrations in 73 bumblebees (43 controls, 20 exposed negative, 10 exposed positive; Table S1). Overall, 71.23% of the bumblebees responded to sucrose (Figures 1B and C). This percentage was similar across the three infection statuses (X^2^ = 2.236, df = 2, p = 0.326). Bumblebees of the three infection statuses responded similarly to the lowest sucrose concentration (Figure 1 B; GLMM, X^2^ = 0.920, df= 2, p = 0.631), reached similar gustatory scores (Figure 1C; GLMM, X^2^ = 1.686, df = 2, p = 0.430) and similar final level of response to the highest sucrose concentration (Figure 1B; GLMM, X^2^ = 0.217, df = 2, p = 0.897). Therefore, exposure to *N. ceranae* affected neither the sucrose sensitivity nor the feeding motivation of bumblebees.

### 3.1. Parasite exposure reduced absolute learning but not memory

We analysed absolute conditioning in 420 bumblebees (141 controls, 228 exposed negative, 51 exposed positive; Table S2). The proportion of bumblebees showing a conditioned response increased over trials in the three infection statuses (Figure 2A). Exposed bumblebees (either negative or positive) had significantly lower acquisition scores (Figure 2B; GLMM, status: X^2^ = 31.558, df = 2, p < 0.001) and learning scores (Figure 2C; GLMM, status, X^2^ = 16.037, df = 2, p < 0.001) than controls. Both acquisition and learning scores were similar for negative and positive exposed bumblebees (Tukey p > 0.05; Table S4).

**Figure 2.**
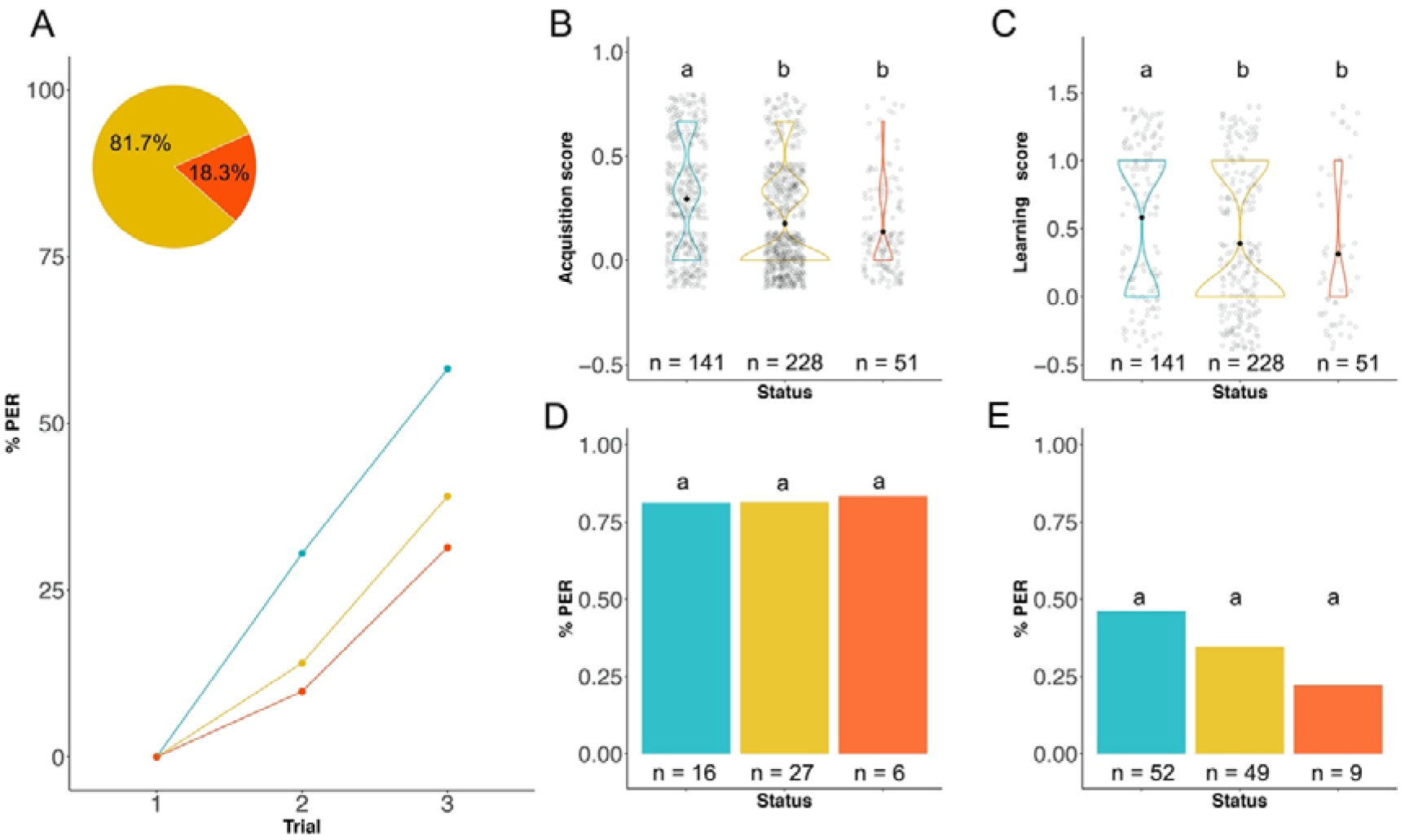
Absolute learning and memory. **A)** Learning curves show the proportion of control (blue), exposed negative (yellow) and exposed positive (red) bumblebees extending their proboscis to the odour during conditioning. Pie chart shows the percentage of negative and positive exposed bumblebees that finished the conditioning (N total = 279). **B-C)** Violin plots for acquisition score (B; sum of correct responses divided by the number of trials for each bumblebee) and learning score (C; correct response at the last trial). Black diamonds represent the mean score for each infection status. White dots represent the score of each individual. **D)** Short-term memory. Bar plots show the proportion of bumblebees responding to odour 1 h after training. **E)** Long-term memory. Bar plots show the proportion of bumblebees responding to odour 24 h after training. Letters above violin plots and bar plots represent significant differences between status (GLMM; p < 0.05). n is the sample size.

We analysed short-term and long-term memory formation of efficient bumblebees that responded to odour at least once in any of the last two last trials (41.52% in short-term memory and 50% in long-term memory; Table S2). A high proportion of the bumblebees (82.02%±0.011, mean±SE) remembered the rewarded odour 1h after training. This proportion was similar in the three infection statuses (GLMM, status: X^2^ = 0.013, df = 2, p = 0.993), indicating that *N. ceranae* did not affect short-term memory (Figure 2D). A much lower proportion of the bumblebees (34.35%±0.06, mean ±SE) remembered the rewarded odour after 24 h. Exposed bumblebees (either negative or positive) showed a lower proportion of correct responses than controls, although this was not significantly different (Figure 2E; GLMM, status: X^2^ = 2.265, df = 2, p = 0.322), presumably due to the low amount of exposed positive individuals (9 bumblebees).

### 3.2. Parasite exposure reduced differential and reversal learning

We analysed differential learning in 125 bumblebees (64 controls, 41 exposed negative, 20 exposed positive; Table S3). Bumblebees in the three infection statuses increased their response to A+ over trials, but not to B- (Figure 3A), indicating that they were equally able to discriminate the odours. However, parasite exposure affected acquisition scores (GLMM, status, X^2^ = 22.039, df = 2, p < 0.001; Figure 3B). Exposed bumblebees (positive or negative) showed similar acquisition scores (Tukey test: p > 0.05; Table 1) and these scores were significantly lower than the scores of controls (Tukey test: p < 0.05; Table 1). Parasite exposure also affected the final level of learning (GLMM, status, X^2^ = 11.168, df = 2, p = 0.003; Figure 2C). Exposed positive bumblebees had the lowest learning score. Their score was significantly lower than that of controls (Tukey test: p = 0.002; Table 1) but similar to that exposed negative bumblebees (Tukey test: p = 0.133; Table 1). Thus overall, exposure to *N. ceranae* reduced differential learning performances.

**Figure 3.**
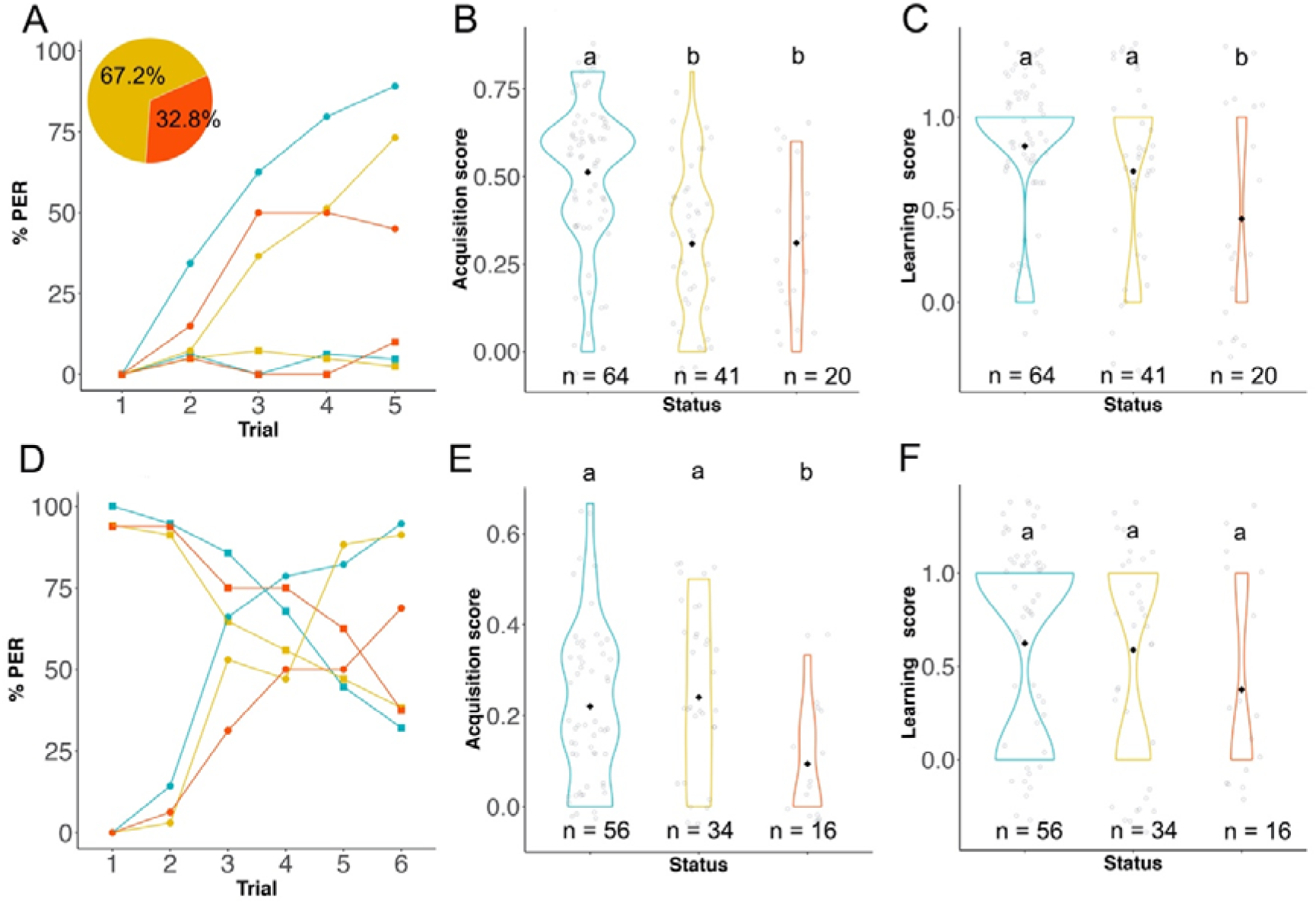
Reversal learning. **A-C)** Differential learning phase. **A)** Percentage of PER responses to rewarded (A+, circle) and unrewarded (B-, square) odours by control (blue), exposed negative (yellow) and exposed positive (red) bumblebees. Pie chart shows the percentage of exposed negative and positive bumblebees that finished conditioning (n total = 67). Violin plots of **B)** acquisition scores (i.e. sum of the correct responses divided by the number of trials for each bee) and **C)** learning scores (i.e. performance of bumblebees at last trial) of bumblebees with different infection statuses. Black diamonds represent the mean score of each status. White dots are the scores for each individual. **D-F)** Reversal learning phase. **D)** Curves show the increase in the percentage of PER response to B+ over A- over trials. **E)** Acquisition score. **F)** learning scores. Letters above violin plots show significant differences between infection statuses in the acquisition and learning scores (GLMM, p < 0.05). n is the sample size.

We analysed reversal learning in 108 bumblebees that finished the differential phase, as they responded to A- in any of the two first presentations of this odour during reversal phase (56 controls, 36 exposed negative, 16 exposed positive; Table S3). All bumblebees reduced their response to A- in favour of B+ over trials (Figure 3D). However, bumblebees with different infection statuses reached significantly different acquisition scores (Figure 3E; GLMM, status, X^2^ = 6.042, df = 2, p = 0.048). Exposed positive bumblebees showed slightly lower acquisition scores than controls (Tukey test: p = 0.049; Table 1) and exposed negative bumblebees (Tukey test: p = 0.056, Table 1), suggesting they had a lower ability to reverse the task. By the end of training, bumblebees of all infection statuses reversed the task, as shown by the lack of difference between their learning scores (Binomial GLMM, status: X^2^ = 3.073, df = 2, p = 0.215). Thus overall, exposure to *N. ceranae* also impaired the reversal phase of reversal learning. We found no effect of *N. ceranae* exposure on either phase of reversal learning when bumblebees were tested at 2 days post exposure, suggesting that stress due to parasite exposure or parasite infection requires longer time to be established (Text S1).

## 4. Discussion

Bees are exposed to a number of parasites that can affect cognitive abilities supporting crucial behaviour (Koch et al., 2017; Schmid-Hempel, 2013). Previous studies exploring the effect of *N. ceranae* on absolute olfactory learning and memory in bees reported contrasting results, presumably because of differences in conditioning protocols and infection rates across studies (Bell et al., 2020; Charbonneau et al., 2016; Gage et al., 2018; Piiroinen et al., 2016; Piiroinen and Goulson, 2016). Here, we ran a suite of standard olfactory cognitive assays showing that feeding bumblebees spores of this parasite consistently impairs different types of olfactory learning but not memory seven days after exposure. This suggests that the influence of *N. ceranae* exposure on bumblebee cognition is not specific to a particular brain area.

Exposure to *N. ceranae* in food clearly impaired the ability of bumblebees to associate an odour with a reward (absolute learning), discriminate two odours (differential learning), and learn an opposite association (reversal learning). These are fundamental cognitive operations a bee must display to efficiently forage on flowers (Giurfa and Sandoz, 2012). This result agrees with a previous study reporting a reduced absolute learning in *N. ceranae* exposed bumblebees (Piiroinen and Goulson, 2016). Here we provide evidence that these reduced cognitive performances were not due to an effect of the parasite on odour perception or motivation to feed, and give insight on where the parasite might act in the nervous system. In bees, sucrose perception through the antennae, olfactory absolute learning, differential learning and reversal learning all require processing of olfactory information through the antennal lobes (Giurfa and Sandoz, 2012) while reversal learning also requires functional mushroom bodies (Boitard et al., 2015). The fact that all types of learning were impaired and that sucrose sensitivity was not, indicates that *N. ceranae* did not specifically target the antennal lobes or the mushroom bodies. Rather, it likely impacted the learning processes in general.

By contrast, we found no evidence that *N. ceranae* influenced memory. During training animals learn and form short-term memories that are later consolidated and transformed into stable long-term memories (Menzel and Muller, 1996) after protein synthesis (Menzel, 2001). In our experiments, *N. ceranae* neither impaired short-term nor long-term memory. Nearly all exposed bumblebees (either positive or negative in PCR) remembered the association of the odour with the reward 1 h after training. However, they showed slightly lower performance than controls at 24 h. This tendency was not significant and may result from low sample size as only 29.44% of the efficient exposed bumblebees were eligible to analyse their memory (i.e. efficient bumblebees, with at least one conditioned response during learning, that responded to sucrose after memory test).

It has recently been questioned whether bumblebees are natural hosts of *N. ceranae* based on the lack of evidence of parasitic forms inside host cells (Gisder et al., 2020). Several studies have nevertheless reported *N. ceranae* in wild bumblebees at low (e.g. 4.76%; (Sinpoo, 2018)) and high prevalence (e.g. 72%; (Arbulo et al., 2015)). Whether or not bumblebees are suitable hosts for *N. ceranae* replication is confirmed in future studies, our results imply that bumblebees are strongly impacted by an acute exposure to the parasite. Such exposure may be extremely frequent in nature due to the high prevalence of *N. ceranae* in honey bees (Runckel et al., 2011)) that contaminate flowers with spores through physical contact or in their faeces (Graystock et al., 2015). Our protocol of parasite exposure significantly increased the infection rate of bumblebees to 28% in comparison to previous studies (Piiroinen et al., 2016; Piiroinen and Goulson, 2016), which allowed the evaluation of cognitive traits in bumblebees in the three infection statuses. Bumblebees that tested positive to *N. ceranae* showed a tendency for lower cognitive performances than negative exposed bumblebees. They reached the lowest learning during the absolute conditioning and barely discriminated odours, suggesting that infection may interfere with some aspects of cognition, however further experiments are needed to tackle this specific question.

Through which mechanism may the parasite impair learning? Ingestion of *N. ceranae* spores exerts a stress that can reduce cognition. *N. ceranae* is known to alter the immune system of bees, for example by modulating the expression of antimicrobial peptides (Antúnez et al., 2009; Botías et al., 2020; Sinpoo, 2018). Stimulation of the immune system with non-pathogenic elicitors, as lipopolysaccharides (LPS), was shown to reduce learning abilities in honey bees, that were less able to associate an odour with a reward (Mallon et al., 2003), and bumblebees, that showed lower performances in odour (Mobley and Gegear, 2018) and colour differential learning tasks (Mobley and Gegear, 2018). It is thus possible that the observed effects of *N. ceranae* exposure on bumblebee cognition were caused by an activation of the immune response. In honey bees, infection with *N. ceranae* was shown to downregulate the expression of genes in the brain (Doublet et al., 2016), some of which are linked to olfaction (Badaoui et al., 2017; Doublet et al., 2016), potentially leading to changes in behaviour and cognition. Interestingly, none of these effects were observed at two days post exposure (Text S1). Whether *N. ceranae* triggers the immune response at this time is unknown. In honey bees, the earliest effects described so far were observed after three days (Chaimanee et al., 2012).

Since olfactory learning is essential for foraging, this sublethal effect of *N. ceranae* exposure on bumblebee cognition can compromise colony foraging and development. Commercial and wild bumblebee colonies exhibit physiological and behavioural differences as a result of different selective pressures (Velthuis and Doorn, 2006), and may, therefore, show different susceptibility to parasites. Parasite loads in the field can range from a few to thousands spores (Meana et al., 2010). Here we used a substantially higher spore loads of *N. ceranae* to infect commercially reared bumblebees than the actual infection rates found in wild bumblebees (e.g. 6800 spores per individual (Graystock et al., 2013)). Further studies are thus needed to analyse the effects of different concentrations of *N*. ceranae spores, and their possible interactions with other stressors in the field in order to assess their real impact on wild pollinators.

## 5. Acknowledgements

We thank Philipp Heeb, Cristian Pasquaretta and Maria Eugenia Villar for constructive discussions, Charline Schartier for helping running some experiments, and Martin Giurfa for lending the automated PER conditioning setup.

