## Supplementary material for "The gut parasite *Nosema ceranae* impairs olfactory learning in bumblebees"

### Supplementary Materials

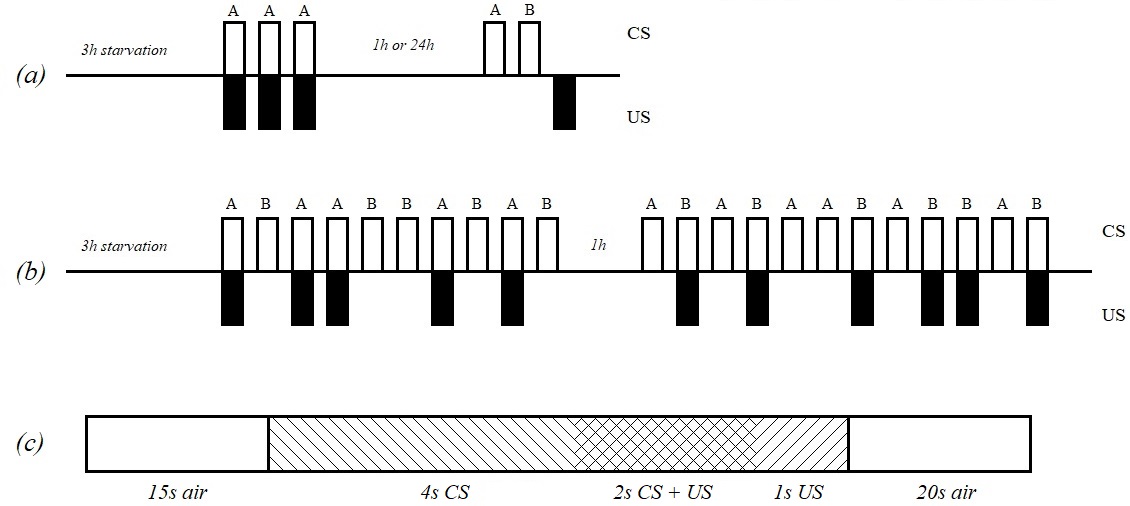

Supplementary Figure S1. Schematic representation of the PER protocols used in cognitive assays using two odorants (A and B). (a) Sequences used in the absolute learning and memory tasks. (b) Sequences used for rewarded and unrewarded trials in the differential and reversal tasks. (c) Sequence of events used in every trial. White bars represent odourless air flow before and after conditioning. Odour (CS) alone is represented by right diagonal lines. Sucrose (US) alone is represented by left diagonal lines. The crossing pattern shows the overlap of CS and US presentation.

Supplementary Table S1. Number of bumblebees tested for sucrose sensitivity at 7 days post exposure.

| *Treatment* | **Control** | **Exposed** |
| --- | --- | --- |
| *Fed* | 48 | 30 |
| *Positive in PCR* | 5 | 10 |
| *Did not finish the test* | 0 | 0 |
| ***Total*** | **43** | **30** |

**Supplementary Table S2**. **Number of bumblebees trained in an absolute learning and tested for short-term memory (STM) and long-term memory (LTM).** Exposed positive bumblebees are shown into brackets. The proportion of bumblebees that failed to respond to the US in at least 1 trial during conditioning (inefficient bumblebees) was similar in control (69.13%) and exposed (69.40%) bumblebees. Only bumblebees that showed a positive conditioning (i.e. conditioned bumblebees) during the learning phase were kept for memory tests. Mortality within 24 h before LTM was 7.6%. Near 18% of exposed bumblebees and 9.6% of control were not motivated to accomplish the memory test (us.memory = 0; Chi.square X^2^ =1.187, df = 1, p-value = 0.279).

| **Cognitive test** | **Bumblebee description** | **Control** | **Exposed (+positive)** |
| --- | --- | --- | --- |
| Absolute learning | *Selected for training* | 234 | 411 |
|  | *Died during training* | 4 | 9 |
|  | *Escaped* | 0 | 1 |
|  | *Not efficient* | 71 | 122 |
|  | *PCR positive* | 18 | 51 |
|  | **Efficient** | **141** | **228 (+51)** |
| Short-term memory (STM) | *Selected for STM* | 36 | 67(+15) |
|  | *Conditioned* | 16 | 27(+6) |
|  | *Died before test* | 0 | 0 |
|  | *us.memory = 0* | 0 | 0 |
|  | **Total to analyse** | **16** | **27 (+6)** |
| Long-term memory (LTM) | *Selected for LTM* | 105 | 161(+36) |
|  | *Conditioned* | 72 | 68(+11) |
|  | *Died before test* | 15 | 7 (+1) |
|  | *us.memory = 0* | 5 | 12(+1) |
|  | *Response to NoD* | 1 | 2 (+0) |
|  | **Total to analyse** | **52** | **49(+9)** |

Supplementary Table S3. Number of bumblebees analysed in reversal learning. Exposed positive bumblebees are shown into brackets. 16.57% of the bumblebees died during the differential phase, 11.25% were not motivated (i.e. did not respond to US in two or more trials), and 4.97% extended its proboscis to the odour at its first presentation. 11.94% of the bumblebees that finished the differential phase died during the reversal phase. Only four bumblebees failed to respond to A- in any of the first two trials with this odour during the reversal phase). We discarded them from the analyses.

|  | ***Control*** | ***Exposed*** |
| --- | --- | --- |
| *Selected for training* | 85 | 96 |
| *Died during differential phase* | 12 | 18 |
| *Unmotivated* | 6 | 11 |
| *Response to odour at first presentation* | 3 | 6 |
| *Positive PCR* | 0 | 20 |
| **Total number of bees used to analyse differential learning** | **64** | **41 (+20)** |
| *Died in reversal* | 6 | 6 (+3) |
| *Failed response to A-* | 2 | 1 (+1) |
| **Total number of bees used to analyse reversal learning** | **56** | **34 (+16)** |

Supplementary Table S4. Tukey Pairwise comparisons for absolute and reversal learning. Significant results (P<0.001) are highlighted in bold.

| Test |  | Contrast | z | P |
| --- | --- | --- | --- | --- |
| Absolute learning | Acquisition score | **control – exposed negative** | **-5.140** | **<0.001** |
|  |  | **control – exposed positive** | **-4.378** | **<0.001** |
|  |  | exposed negative – exposed positive | -1.017 | 0.56 |
|  | Learning score | **control – exposed negative** | **-3.513** | **0.001** |
|  |  | **control – exposed positive** | **-3.174** | **0.004** |
|  |  | exposed negative – exposed positive | -0.996 | 0.572 |
| Reversal learning – differential phase | Acquisition score | **control – exposed negative** | **-4.251** | **<0.001** |
|  |  | **control – exposed positive** | **-3.211** | **0.003** |
|  |  | exposed negative – exposed positive | 0.106 | 0.993 |
|  | Learning score | control – exposed negative | -1.653 | 0.222 |
|  |  | **control – exposed positive** | **-3.323** | **0.002** |
|  |  | exposed negative – exposed positive | -1.981 | 0.133 |
| Reversal learning – reversal phase | Acquisition score | control – exposed negative | 0.152 | 0.987 |
|  |  | **control – exposed positive** | **-2.337** | **0.049** |
|  |  | exposed negative – exposed positive | -2.279 | 0.056 |

**Supplementary Text S1.** **Reversal learning at two days post exposure.**

At 2 days post exposure we analysed differential learning in 104 bumblebees (see details in Supplementary Table S4). *N. ceranae* did not affect differential learning nor reversal learning (Supplementary Figure S2). During the differential learning phase, bumblebees in all infection statuses increased their response to A+, but not to B-, over trials (Figure S3A), indicating their ability to discriminate between odours. Performance was similar in the three infection statuses that reached similar acquisition score (Figure S3B, GLMM, status: X^2^ = 1.235, df = 2, p = 0.539) and learning score (Figure S3C; Binomial GLMM, status: X^2^ = 1.185, df = 2, p = 0.552) by the end of the test. During the reversal learning phase, bumblebees reversed previous contingency as shown by the decrease in the proportion of responses to A- in favour to B+ along trials (Figure S3D). Neither acquisition (Figure S3E; GLMM, status: X = 2.83, df = 2, p = 0.242) nor learning scores (Figure S3F; Binomial GLMM, status: X = 2.053, df = 2, p = 0.358) were affected by parasite exposure, all groups being able to reverse the task.

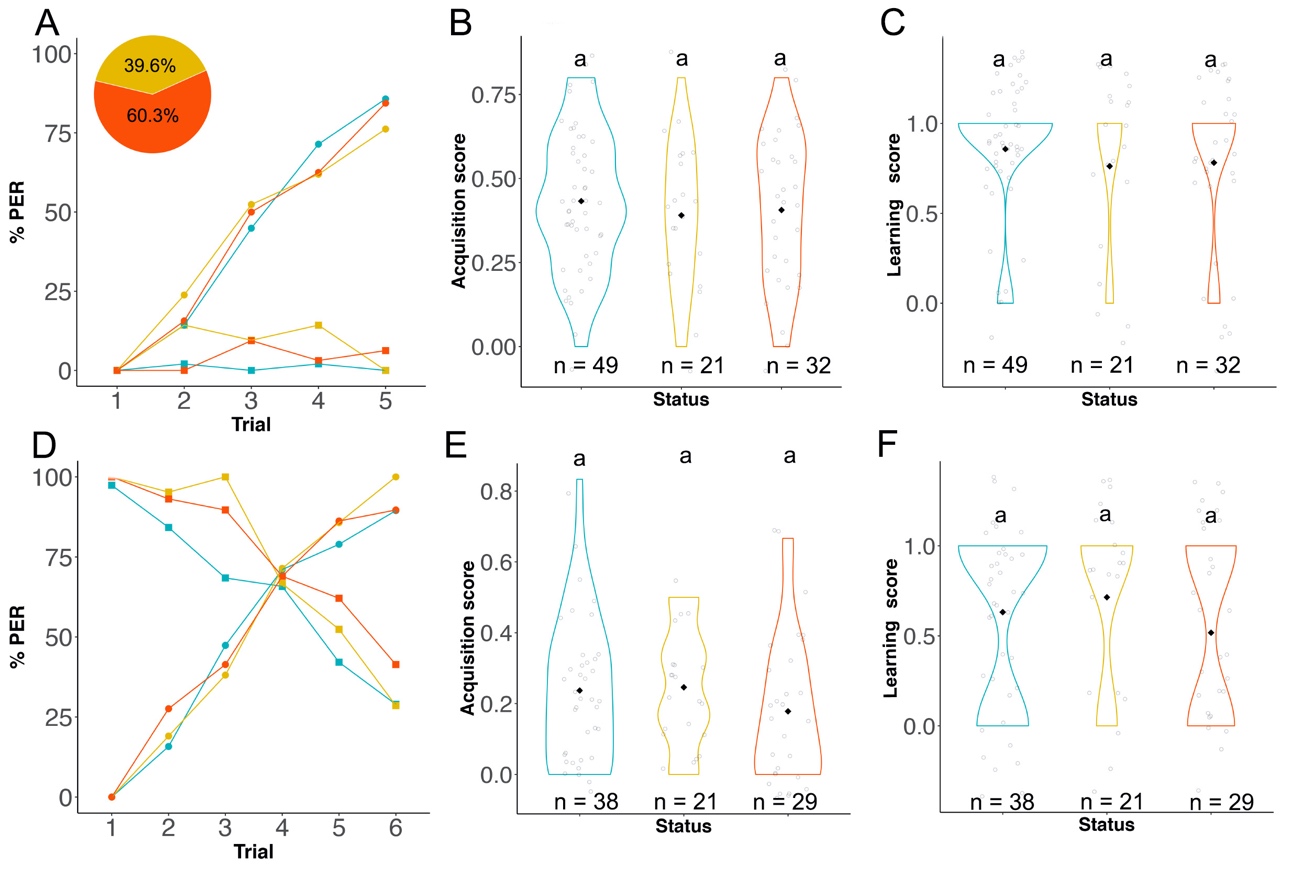

**Supplementary Figure S2*.* Reversal learning at 2 days after exposure.** **A-C)** Differential learning phase. **A)** Curve showing the percentage of PER responses to rewarded (A+, circle) and unrewarded (B-, square) odours by control (blue), exposed negative (yellow) and exposed positive (red) bumblebees. Pie chart shows the percentage of exposed bumblebees that finished the phase testing positive (red) and negative (yellow) to *N. ceranae* in a PCR. Violin plots of **B)** acquisition scores (i.e. sum of the correct responses divided by the number of trials for each bee) and **C)** learning scores (i.e. performance of bumblebees at last trial) of bumblebees with different status. Black dots represent the mean score of each status. Hollow dots represent the score of each individual. **D-F)** Reversal learning phase. **D)** Curves show the increase in the proportion of PER responses to B+ over A- over trials. **E)** Acquisition and **F)** learning scores during reversal phase. Letters above violin plots show significant differences between status in the acquisition (GLMM, p < 0.05) and learning (Binomial GLMM) scores. n is the sample size.

Supplementary Table S5. Number of bumblebees analysed in reversal learning. Exposed positive bumblebees are shown into brackets. During the differential phase 10.6% of the bumblebees died, 0.75% did not respond to US in at least two trials, and 6.81% responded to CS at its first presentation. We discarded them from the analyses. We also discarded six control bumblebees that tested positive to *N. ceranae* in the PCR. In the reversal phase five bumblebees did not fit the selection criteria (i.e. PER to A- in any of the first two trials with this odour during the reversal phase) and were not taken into account for the analyses. 8.82% of bumblebees died during this phase.

|  | *Control* | *Exposed* |
| --- | --- | --- |
| Fed | 65 | 67 |
| *Died in differential* | 5 | 9 |
| *Unmotivated* | 1 | 0 |
| *Response to odour at first presentation* | 4 | 5 |
| *Positive PCR* | 6 | 32 |
| Total Differential | 49 | 21 (+32) |
| *Died in reversal* | 7 | 0 (+2) |
| *Failed selection criteria* | 4 | 0 (+1) |
| Total Reversal | 38 | 21(+28) |
